# The FERONIA receptor kinase is required for high humidity responses in Arabidopsis

**DOI:** 10.64898/2026.04.24.720662

**Authors:** Comzit Opachaloemphan, Richard Hilleary, Jingyuan Zhang, Sooyeon Park, Nan Wu, Chi Kuan, Kinya Nomura, Sheng Yang He

## Abstract

High humidity greatly influences plant growth and development and triggers adaptive physiological responses such as leaf hyponasty (elongation of leaf petiole and upward leaf movement). Cyclic Nucleotide-Gated Ion Channels 2 and 4 (CNGC2/4), Calmodulin Binding Transcription Activators 2 and 3 (CAMTA2/3)-dependent transcription, and ethylene signaling acting through EIN2 and EIN3/EIL1 are required for high humidity response in Arabidopsis. However, upstream regulators controlling these steps remain unknown. Here, we show that the receptor-like kinase FERONIA and its co-receptor LORELEI-LIKE GPI-ANCHORED PROTEIN1 (LLG1) are required for a large portion of high humidity-associated Arabidopsis transcriptomic changes, including *CNGC2*, *CAMTA*-regulated genes, and cell wall remodeling genes, and for high humidity-induced leaf hyponasty. High humidity triggers petiole-localized Ca²⁺ waves that precede hyponastic leaf movement. The petiole-localized Ca^2+^ signals were also significantly altered in the *fer-4* mutant. Thus, the FERONIA receptor kinase is a critical early regulator of plant responses to high humidity.

## INTRODUCTION

To survive, plants rely on adaptive responses enabling their survival and resilience in dynamic environments^<u>1,2</u>^. Among these environmental variables, air humidity exerts a strong influence on plant biology. Similarly to thermomorphogenesis and shade avoidance, high humidity promotes petiole elongation and upward leaf movement (hyponasty), which are thought to improve cooling, airflow, and/or photosynthetic positioning^<u>3–6</u>^. High humidity also favors microbial colonization by creating an aqueous leaf apoplast^<u>1,7,8</u>^. Despite the prominent effects of high humidity on plant biology and plant interaction with microbes, the molecular details of plant response to high humidity have only emerged in the past decade^<u>4,8–14</u>^.

High humidity influences multiple hormone pathways, including abscisic acid (ABA), jasmonic acid (JA), salicylic acid (SA), and ethylene^<u>9,12–14</u>^. A recent study showed that high humidity triggers CNGC2/4-mediated Ca²⁺ influx and CAMTA2/3-dependent transcription, including *CYP707A3* involved in ABA catabolism^<u>15</u>^. These findings underscore that high humidity reprograms hormonal and calcium signaling networks. However, how plants sense atmospheric high humidity and initiate these signaling cascades remains unclear.

Arabidopsis encodes more than 600 receptor-like kinases (RLKs) that perceive diverse extracellular cues^<u>16–18</u>^ and regulate processes ranging from development and hormone perception to pathogen detection and cell wall integrity^<u>18–20</u>^. Among RLKs, FERONIA contains a malectin-like ectodomain that binds RAPID ALKALINIZATION FACTORS (RALFs) and pectin *in vitro,* and sensing cell wall integrity *in vivo*^<u>21–25</u>^. Here, we identify FERONIA as an important early mediator of high-humidity responses in Arabidopsis. Through transcriptomics, mutant screening, and whole-plant Ca²⁺ imaging, we show that FERONIA is required for rapid petiole-localized Ca²⁺ waves, induction of high humidity-associated transcriptome, including *CNGC2* and a suite of hormone and cell wall-remodeling genes, and hyponastic growth.

## RESULTS

### Whole leaf transcriptomic analysis of Arabidopsis response to high humidity

To begin dissecting the Arabidopsis transcriptomic response to high humidity, we conducted whole-leaf RNA-seq analysis of 4-week-old Col-0 plants exposed to high humidity (RH > 95%) for 15 minutes, 60 minutes, or 24 hours to capture early and late timepoint transcriptomic responses (Figure 1A). At 15-minutes, 732 differentially expressed genes (DEGs) were identified (565 up- and 167 down-regulated DEGs compared to no high humidity exposure controls). The most pronounced changes were observed at 60 minutes with 4,508 up- and 4,295 down-regulated DEGs, whereas 2,180 up- and 2,209 down-regulated DEGs were found at 24-hour time point (Figure 1A; Figure S1; Table S1).

**Figure 1.**
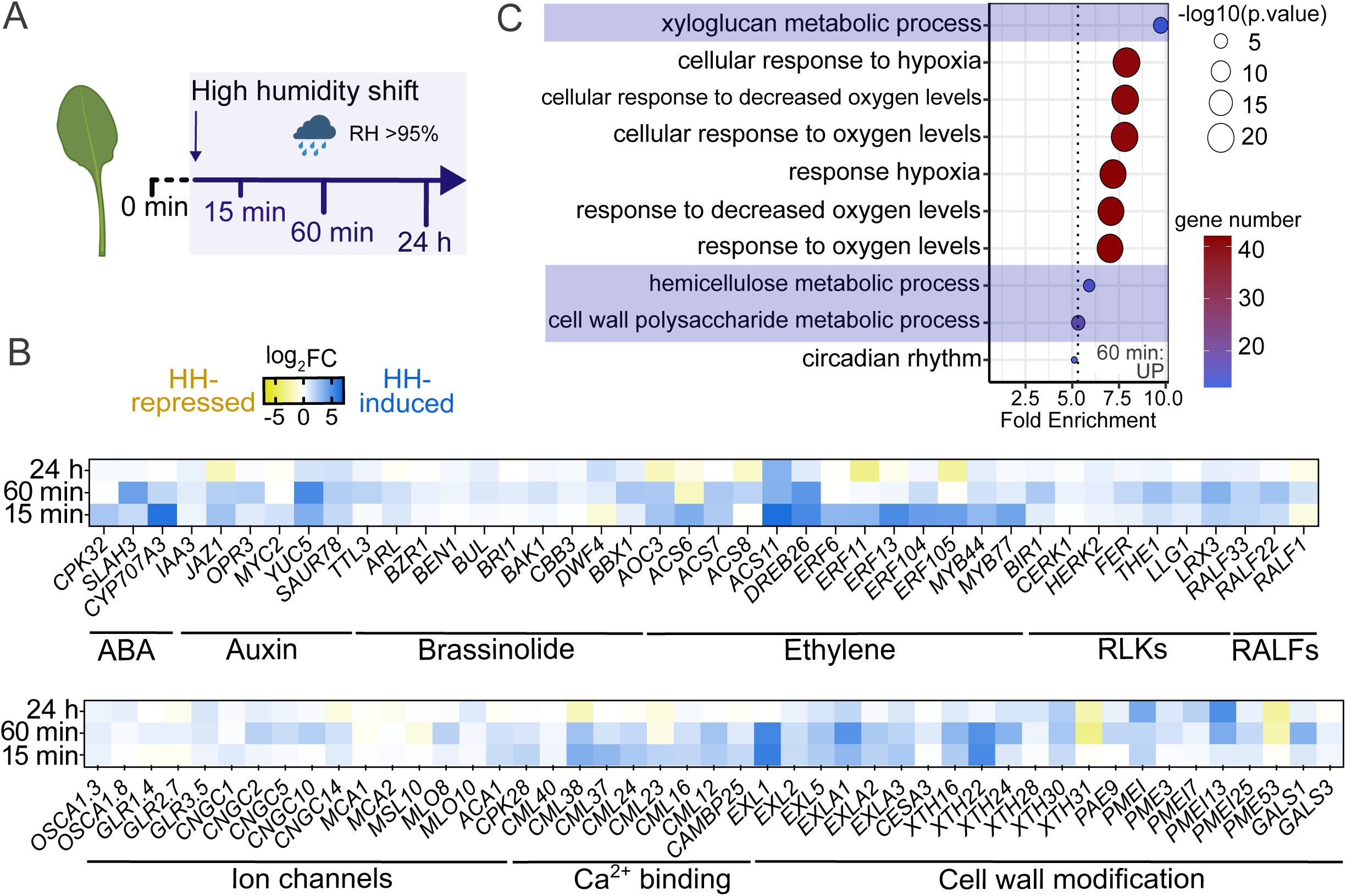
Rapid high humidity-responsive changes in cell wall gene expression. (A) Schematic overview of RNAseq experimental setup and leaf sample collection at four time points. (B) A heatmap shows the expression profiles of selected differentially expressed genes in different categories across the 15-minute, 60-minute and 24-hour timepoints. Color scale represents log_2_ fold change (after high humidity treatments vs. before treatment). Blue indicates high-humidity induction and yellow indicates repression (p-adjusted < 0.05). (C) Gene Ontology (GO) enrichment analysis of high humidity-induced genes at the 15-minute timepoint. Biological processes significantly enriched genes are shown and sorted by fold enrichment.

Gene ontology (GO) enrichment analysis of up-regulated DEGs at 15 and 60 minutes revealed strong signatures of hypoxia responses, along with plant cell wall remodeling (Figures 1B and 1C; Figure S1C). Detailed inspection of induced genes, however, revealed induction of additional genes encoding FERONIA and other RLKs, RALFs, ion/calcium-permeable channels, and hormone-associated genes linked to abscisic acid (ABA), auxin, ethylene and brassinosteroid (BR) pathways (Figure 1B; Table S1). Notably, *CYP707A3*, an ABA catabolic gene and an early high-humidity marker activated by CAMTA2/3 downstream of Ca²⁺ influx^<u>15</u>^, was strongly induced in Col-0. Likewise, several Ca²⁺-associated transcripts (e.g., *CNGC2*, *ACA1, CAMBP25, CMLs, CPK28*) and ethylene signaling genes including *ACS/ERF*-family members^<u>26</u>^ were induced (Figure 1B).

Pectin is a major hydrophilic component of the cell wall, and its biological activity is strongly influenced by the degree of methyl esterification^<u>26–30</u>^. We found that high humidity altered the expression of numerous pectin-related genes in Col-0, including those coding for galactan synthases (*GALSs),* galactosyltransferases (*GAUTs*), pectin lyases (*PLs*), pectin acetylesterases (*PAEs*), pectin methylesterases (*PMEs*), and PME inhibitors (*PMEIs*) (Figure 1B). Clustered *PMEI–PME* genes, which encode proteins with an intramolecular PMEI domain that regulates PME activity^<u>30,31</u>^, were robustly induced by high humidity in Col-0. Consistent with the altered expression of pectin-related genes, we found that Col-0 ultimately exhibited increased levels of de-esterified pectin after high humidity treatment, compared to plants under ambient humidity (Figure S1E). Taken together, our temporal transcriptional analysis suggests that high humidity rapidly induces hypoxia, RALF, receptor and hormone signaling pathways, along with cell wall remodeling genes.

### Diminished leaf hyponastic responses to high humidity in the *fer-4* mutant

Guided by the GO enrichment analysis at the early timepoints (15 min and 60 min), we screened a large number of T-DNA mutants covering a range of putative cellular pathways to assess their roles in petiole elongation under high humidity (Table S2). The majority of mutants defective in hypoxia signaling (e.g. *RAP2*, *PBGs* and *PCOs*), ethylene response (e.g. *ERFs*), RLKs (e.g. *THE1*), cell wall pectin modification (e.g. *GALS1/2/3*) and ion channel function (e.g. glutamate receptor-like [*GLRs*], mechanosensitive channel of small conductance-like [*MSLs*] and osmosensitive calcium-permeable channels [*OSCA*s]) exhibited normal petiole elongation in response to high humidity (Table S2; Figure S2). In contrast, the loss-of-function *FERONIA* mutant *fer-4* showed significantly reduced petiole elongation under high humidity (Figures 2A and 2B). To ascertain that the impaired petiole elongation in the *fer-4* mutant is not due to general growth defects, as indicated in its smaller rosette size compared to Col-0 plants, we examined other well characterized growth-impaired, small-size mutants, such as *snc1* (an auto-active nucleotide-binding leucine-rich repeat immune receptor mutant^<u>32</u>^) and *siz1* (E3 SUMO protein ligase mutant^<u>33</u>^), under high humidity. Like Col-0 plants, *snc1* and *siz1* mutants showed comparable relative levels of petiole elongation in response to high humidity, whereas the *fer-4* mutant showed pronounced impairment in high humidity-induced petiole elongation (Figure 2B).

**Figure 2.**
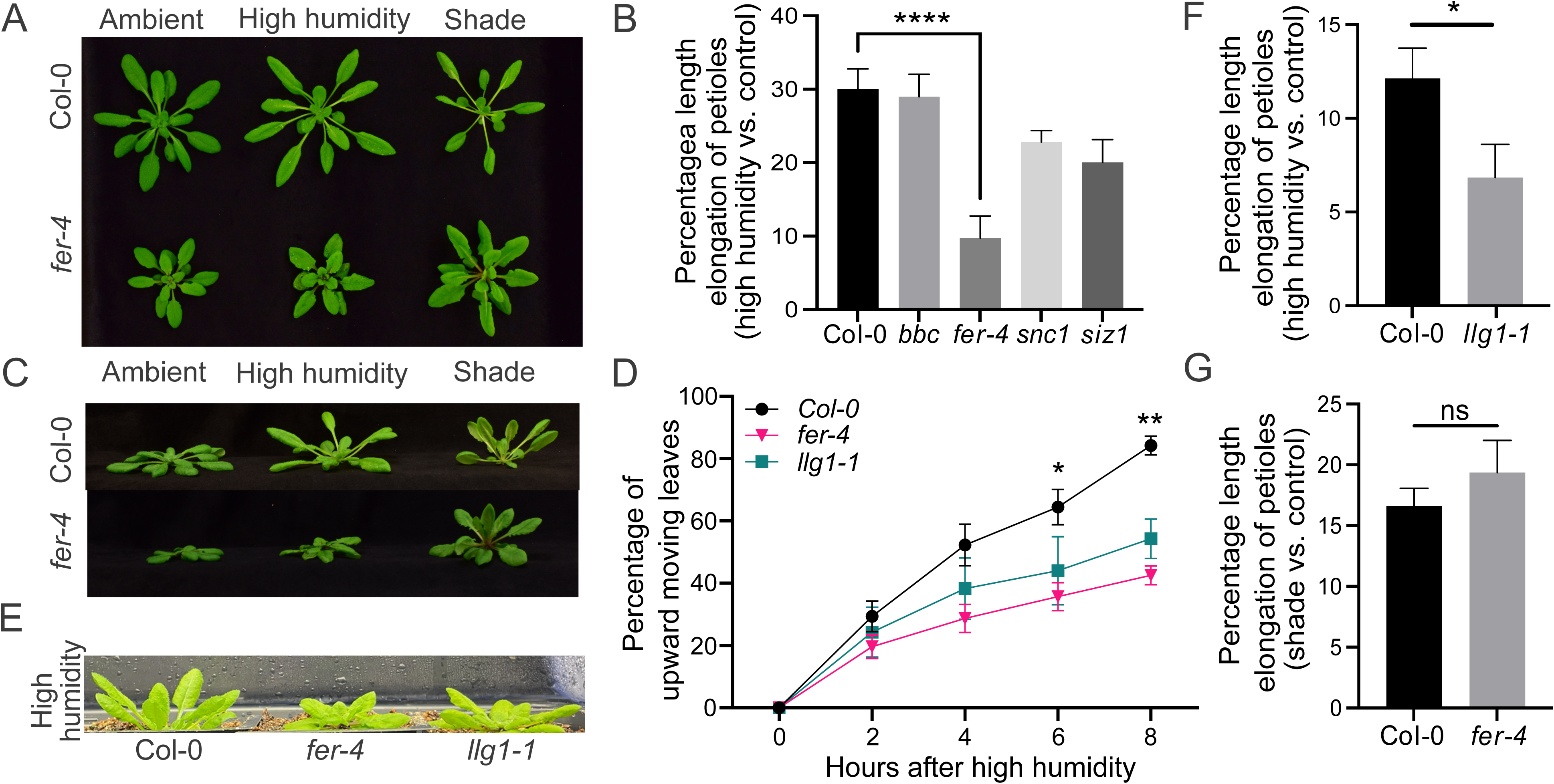
Impaired high humidity responses in the *fer-4* mutant. (A, C) Responses of petiole elongation and upward leaf movement in four-week-old Col-0 and *fer-4* plants after 4 days of high humidity or shade treatments. (B and G) Quantitative plots of petiole-to-leaf length % increase in Col-0 and *fer-4* mutant plants (B) 6 days after plant growth under >95% RH vs. ambient humidity (∼60% RH) condition or (G) 4 days after shade condition. (D) Proportion of upward moving leaves under >95% RH high humidity condition during the first eight hours in Col-0 (black), the *fer-4* mutant (pink) and the *llg1-1* mutant (green). Bars represent mean ± S.E.M; n=4. (E and F) Petiole elongation under >95% RH high humidity condition in Col-0 and the *llg1-1* mutant with a quantitative plot of petiole length % increase six days after ambient and high humidity treatment. Bars represent mean ± S.E.M; n=4. Mann-Whitney U test was used for all experiments. P values were indicated (*p < 0.05, **p < 0.01, ****p < 0.0001, and ^ns^ p > 0.05).

In addition to the petiole elongation assay, which requires a long period of high humidity treatment, we examined leaf movement within the first eight hours of high humidity exposure. Nearly all Col-0 leaves exhibited active and persistent upward movement, whereas *fer-4* leaves showed markedly reduced responsiveness to high humidity, especially leaves in the lower whorl close to the soil (Figure 2D and Video S1). Importantly, we also tested the shade avoidance responses in the *fer-4* mutant and observed normal petiole elongation and upward leaf bending when shaded for 4 days (Figures 2C and 2G). This result shows that FERONIA has a selective impact on high-humidity response rather than causing impairment in leaf hyponasty. These observations showed that FERONIA is involved in facilitating both the early leaf movement response and sustained petiole elongation response to high humidity treatment.

The FERONIA receptor functions with *LORELEI-like GPI-ANCHORED PROTEIN1* (*LLG1*). Interestingly*, LLG1* is also rapidly induced in response to high humidity (Figure 1B). Similar to the *fer*-4 mutant, the *llg1-1* mutant exhibited reduced petiole elongation and leaf movement under high humidity, underscoring the importance of the FER-LLG1 module in high humidity responses (Figures 2E and 2F; Video S1). We also tested whether RALF1, a known FERONIA ligand^<u>34</u>^ induced by high humidity (Figures S2B and S2C), to determine if it contributes to petiole elongation. Treatment Col-0 and *fer-4* with synthetic 20 nM and 1μM RALF1 peptides under both ambient and high humidity conditions did not produce a significant effect on petiole elongation (Figure S2C), suggesting exogenous *RALF1* application alone is not sufficient to drive humidity-dependent petiole elongation.

### FERONIA plays a prominent role in early transcriptomic responses to high humidity

The phenotypic requirement of FERONIA for leaf hyponastic responses suggest that it may be necessary for the transcriptomic responses to high humidity. To investigate the extent of FERONIA involvement in high humidity-induced transcriptomic changes, we performed RNA-seq analysis of Col-0 and *fer-4* plants, at 15 minutes, 60 minutes, 24 hours, and 6 days after exposure to high humidity (Figures S3A, S4 and S5). In addition, we reasoned that tissue-resolved analyses might reveal distinct transcriptional fingerprints involved in FERONIA-mediated high humidity responses, so we sampled leaf blade and petiole tissues separately in this set of RNA-seq experiments.

At the 15-minute timepoint, Col-0 mounted a broad transcriptomic response, as described above for the first set of RNA-seq experiments, whereas the *fer-4* mutant was strongly attenuated (Figures S3-S5). In leaf blades, Col-0 induced 922 genes compared to only 388 in *fer-4* (∼58% loss). In petioles, the effect was even more pronounced: Col-0 induced 423 genes, whereas only 40 responded in *fer-4* (∼91% loss; Figures S3B and S3C). Thus, FERONIA plays a prominent role in configuring the early petiole transcriptome in response to high humidity.

Among FERONIA-dependent genes are ABA catabolism and ethylene signaling genes. For example, *CYP707A3*, an early high-humidity marker gene activated by CAMTA2/3 downstream of Ca²⁺ influx^<u>12,15</u>^, was strongly induced in Col-0 leaf blades and petioles but was essentially unresponsive in the *fer-4* mutant (Figures 3A and 3C). Likewise, several Ca²⁺-associated transcripts (e.g., *CNGC2*, *ACA1, CAMBP25, CMLs, CPK32*) and ethylene signaling genes including *ACS/ERF*-family members showed *FERONIA*-dependent induction in response to high humidity. Some of high humidity-regulated auxin- and BR-associated genes, including *IAA6*, *DWF1* and *BRI1*, were also *FERONIA*-dependent (Figure 3A; Figure S6). The anion channel gene *SLAH3*, a high-humidity marker gene that depends on ethylene signaling^<u>4</u>^, was induced in Col-0 leaf blades and petioles at 60 min but failed to respond in the *fer-4* mutant (Figure 3A; Figure S4E). Similarly, genes associated with cell wall modification were broadly FERONIA-dependent (Figure S4C). Selected arabinogalactan proteins (*AGP2/7/18/22*), *EXPANSIN*- and *EXPANSIN-LIKE*genes (*EXLA1*/2, *EXGT*-*A*1), *XYLOGLUCAN ENDOTRANSGLUCOSYLASE/HYDROLASES* (*XTH9*/*16/17/30*) and *PECTIN*- and *GALACTAN*-*MODIFYING ENZYMES* (*GALS1*, *GAUT4, PME9, PMR6*) were predominantly higher in Col-0 than in *fer-4* across both tissues, with the difference most pronounced at 60 min and in petioles; only a minority of transcripts showed the opposite pattern. The cell wall-related gene *GALS1* was rapidly induced in both leaf blades and petiole of Col-0 but not in the *fer-4* mutant (Figure 3A; Figure S5B). *GALS1* also displayed the strongest FERONIA dependence of this group (petiole, 60 min). The *PECTIN ACETYLESTERASE* gene *PAE9* illustrates this behavior (Figure S4D). *PAE9* was unchanged at 15 min but rose sharply at 60 min in Col-0 leaf blade and petiole, and returned to baseline by 6 d. This induction was strongly attenuated in the *fer-4* mutant in leaf blades and petioles at both 60 min and 24 h, whereas *fer-4* transcript levels fell below ambient by 6 d.

**Figure 3.**
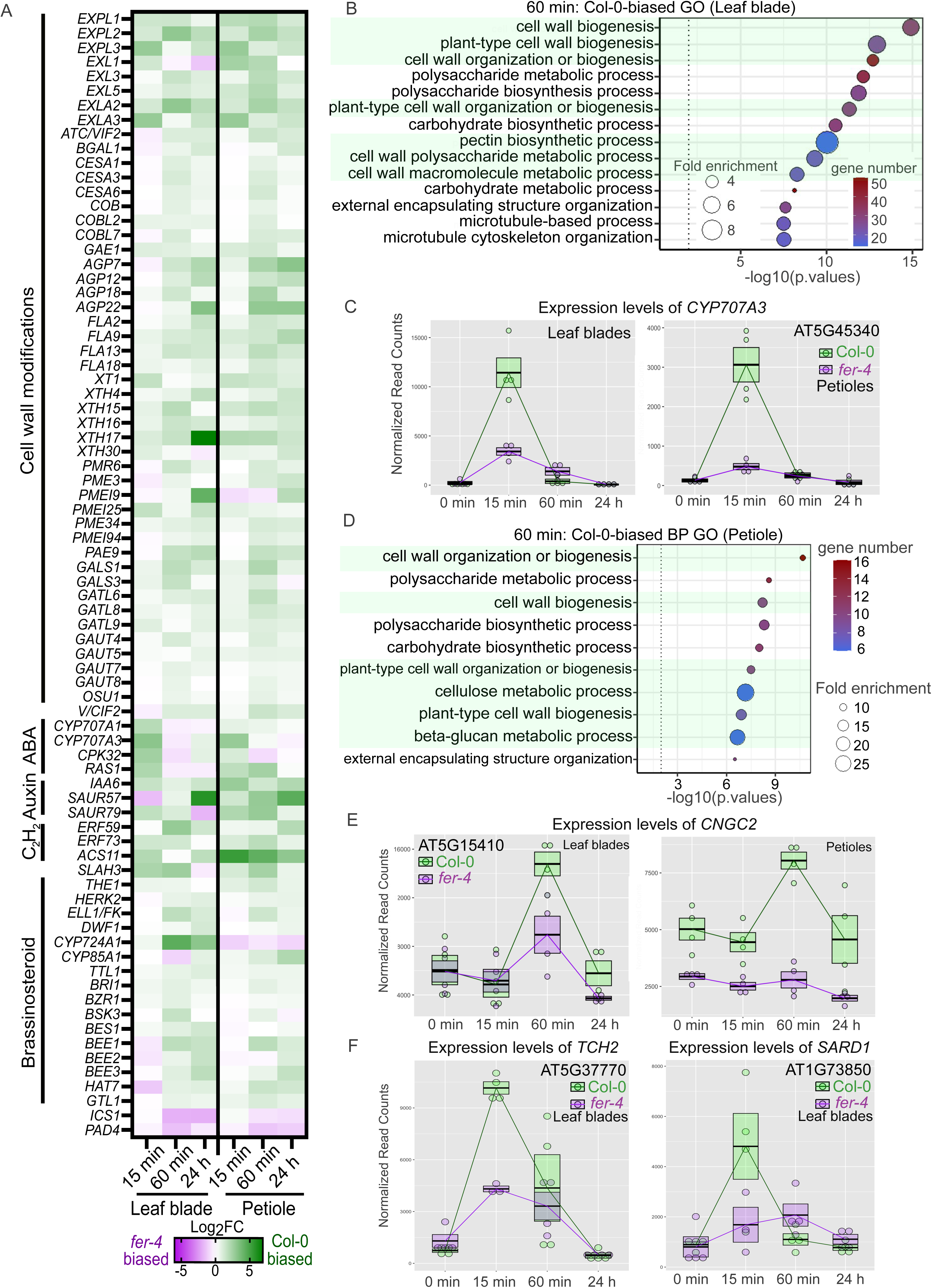
High humidity-responsive cell wall gene expression in leaf blade and petiole tissues of Col-0 and *fer-4*. (A) A heatmap of high humidity-responsive genes in Col-0 but non-responsive in *fer-4* shows log_2_ fold change (Col-0 vs. the *fer-4* mutant). Green color represents highly up-regulated genes in Col-0 (Col-0-biased genes). Purple displays down-regulated genes in Col-0 and higher expressed in the *fer-4* mutant (*fer-4*-biased genes). (B) Biological process (BP) GO enrichment analysis of high humidity-responsive genes uniquely found in Col-0 leaf blades at the 60-minute timepoint. Common biological process GO terms are highlighted in green and sorted by p-values. (C) RNAseq plots of *CYP707A3* gene in the leaf blades and petioles of Col-0 and *fer-4* at 0 minute, 15 minute, 60 minute and 24 hour after >95% RH exposures. (D) BP GO enrichment analysis of high humidity-responsive genes uniquely found in Col-0 petioles at the 60-minute timepoint. (E-F) RNAseq plots of *CNGC2, TCH2 and SARD1* genes in the leaf blades and/or petioles of Col-0 and *fer-4* at 0 minute, 15 minute, 60 minute, and 24 hour after >95% RH exposures and after 6 days at ambient (Amb) and high humidity (HH) conditions. Boxes represent mean and the 25^th^-75^th^ percentiles; n=4.

Finally, *CNGC2* expression at 60 minutes was FERONIA-dependent in petioles but only partially reduced in blades, whereas *CNGC4* and *CAMTA2/3* expression appeared FERONIA-independent (Figure 3E; Figure S6). Because CAMTA2/3 are required for high humidity-induced *CYP707A3* expression and for leaf hyponasty^<u>9,12,15</u>^, we asked whether the expression of known CAMTA-dependent genes requires FERONIA. All twelve gene-tissue combinations of the reported CAMTA-dependent gene set (e.g., *CYP707A3*, *CPKs*, *CAM2*, *EXLA1, TCH2*, *TCH3*, *CBF2*, *SARD1*)^<u>15,35–37</u>^ showed reduced induction in the *fer-4* mutant (Figure 3A; Figure S6). These results suggest that FERONIA is required for the induction of CAMTA2/3-dependent genes independently of *CAMTA2/3* expression levels.

### Ca^2+^ signaling is impaired in the *fer-4* mutant in response to high humidity

We were intrigued by the observation that FERONIA is required for induction of several Ca²⁺-associated genes, including *CNGC2*, which has recently been demonstrated to be critical for Arabidopsis response to high humidity^<u>15</u>^. We therefore asked if the *fer-4* mutant exhibited altered Ca^2+^ signaling dynamics in response to high humidity. Our initial whole-plant imaging of wild-type Col-0 plants expressing the R-GECO1 calcium biosensor^<u>27</u>^ revealed cytosolic Ca²⁺ signal, often within 30 minutes, in the leaves upon exposure to high humidity upon shifting plants from ambient humidity (∼60% RH) to high humidity (>95% RH) (Figure 4A). Cytosolic Ca^2+^ signal in response to high humidity was also observed in GCaMP3-expressing *Nicotiana benthamiana and Nicotiana tabacum*, indicating that this response is broadly conserved across taxa (Figure 5; Videos S2-S4).

**Figure 4.**
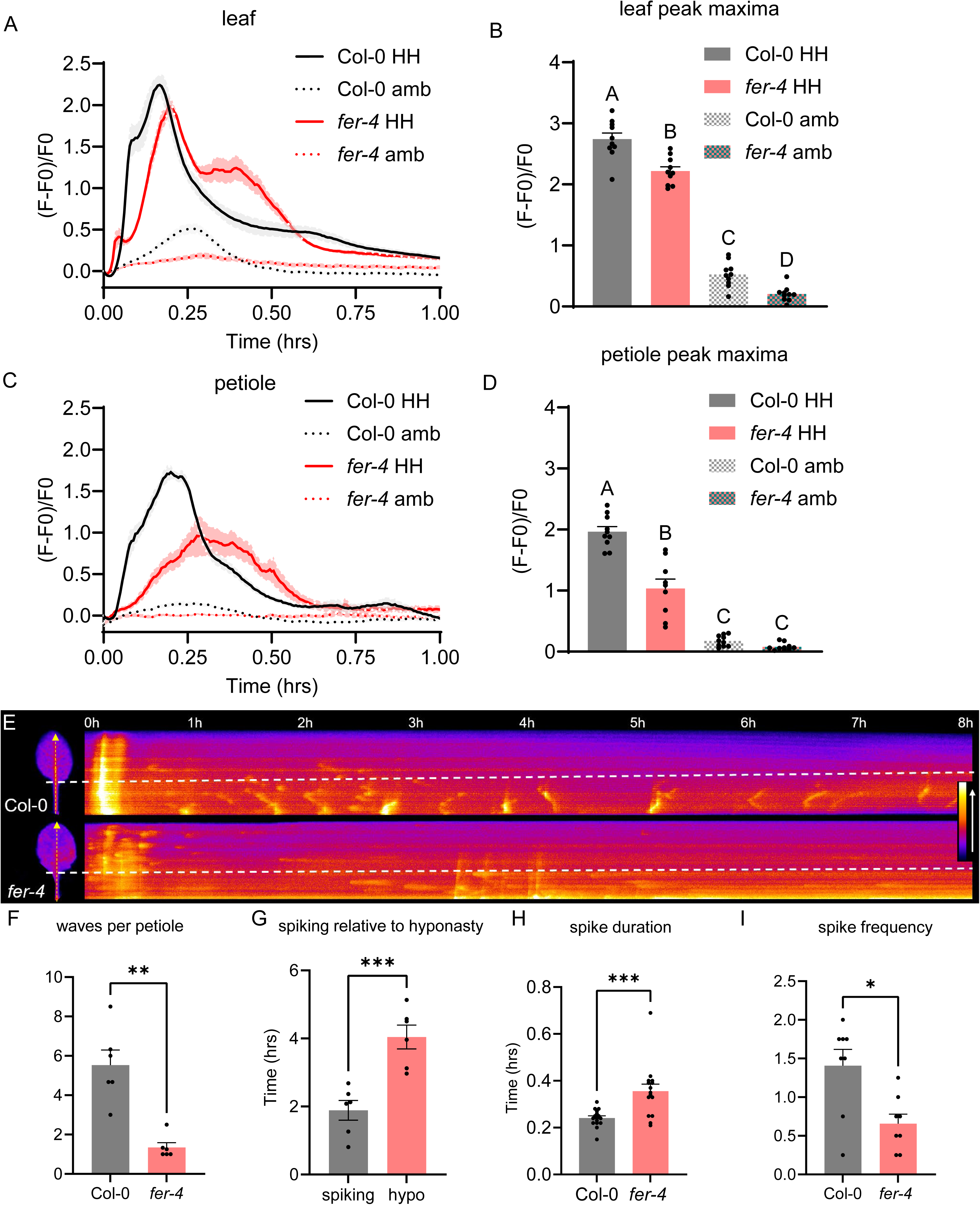
High humidity–induced Ca²⁺ signaling is perturbed in the *fer-*4 mutant. Whole plants expressing the Ca²⁺ biosensor R-GECO1 were shifted from ambient ∼60% RH to >95% RH high humidity. (A-D) Quantification of Ca²⁺ signals in mature leaves (A) and petioles (C) showing mean ΔF over time and peak ΔF over 1 h with peak maxima for leaves (B) and petioles (D) quantified, respectively. (E) Kymographs of rosette leaves and petioles over 8 h highlight spatiotemporal Ca²⁺ waves patterns in Col-0 and *fer-4;* inset color scale shows magenta to orange shifts indicating increased cytosolic Ca^2+^ (yellow arrows = ROIs; dashed lines = leaf blade-petiole border). (F-I) Petiole-localized wave number, spike timing relative to hyponasty, spike duration, and spike per 8 hr. Data are mean ± S.E.M. (n = 12 [A, B] or 6 [E] leaves or petioles from single plants) from one representative experiment. In (B,D), significance was determined by two-way ANOVA (factors were genotype and treatment) followed by Tukey’s post hoc test; different letters indicate significant differences between groups (p<0.05). In (F-I), significance was determined byStudent’s t-test (*p < 0.05, **p < 0.005, ***p < 0.0005).

**Figure 5.**
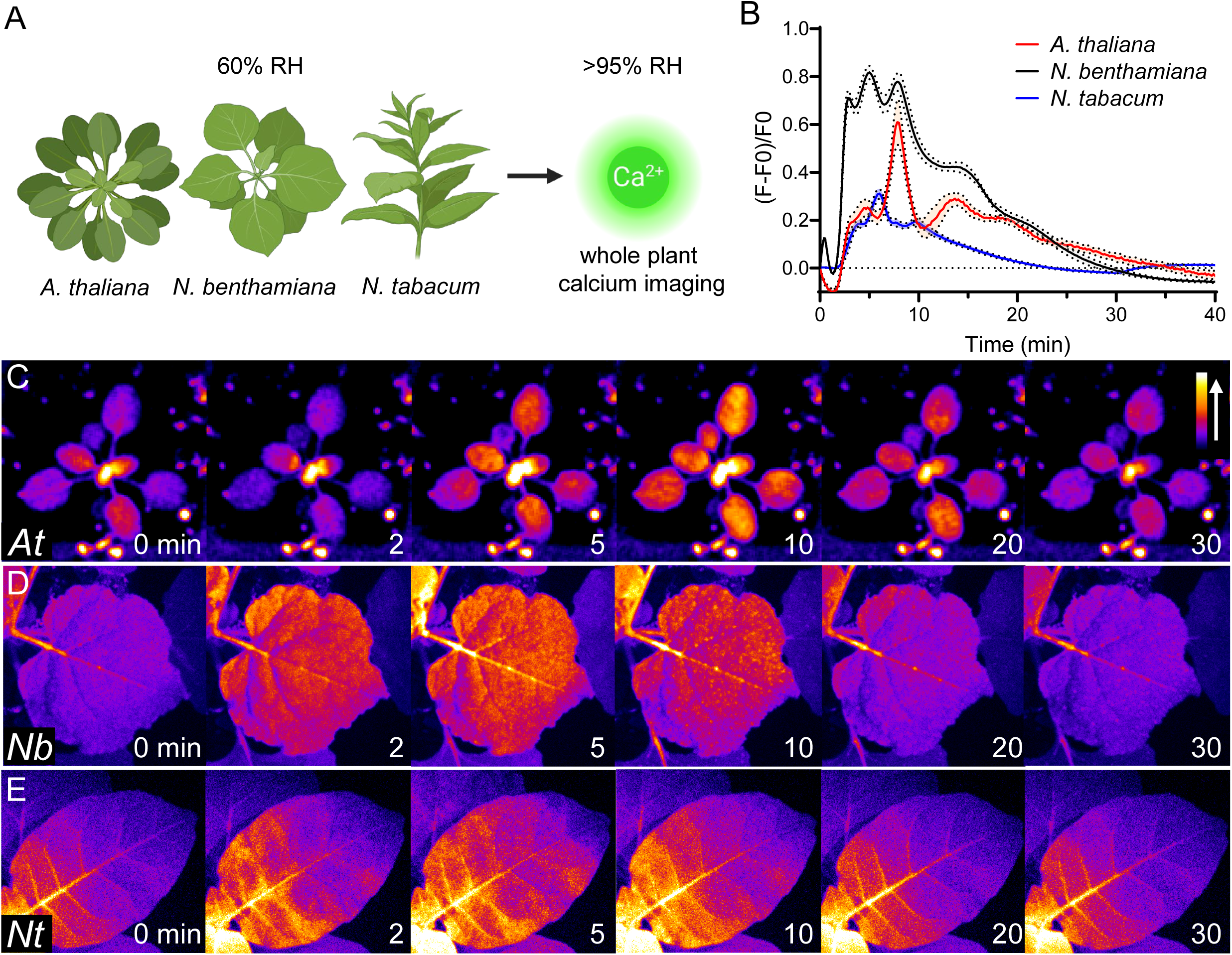
High humidity elicits Ca²⁺ signaling in multiple plant species. Whole plants expressing the Ca²⁺ biosensor GCaMP3 were shifted from ∼60% RH to >95% RH and imaged for Ca²⁺ dynamics. (A) A diagram of experimental setup. (B) Quantification of mean ± S.E.M. Ca²⁺ signals across all leaves (n = 5–7 plants). (C-E) Time-series images showing high-humidity–induced Ca²⁺ signals in *A. thaliana* (C), *N. benthamiana* (D), and *N. tabacum* (E). Color bar shows a shift from magenta to orange indicates increased cytosolic Ca²⁺.

Next, we compared Ca^2+^ responses in Col-0 vs. *fer-4* Arabidopsis leaves. While neither Col-0 nor *fer-4* leaves exhibited an obvious elevation of baseline cytosolic Ca^2+^ at ambient humidity, both Col-0 and *fer-4* genotypes displayed an early cytosolic Ca^2+^ increase (i.e., within 30 minutes) in response to high humidity, with Col-0 showing significantly higher peak signals in both the leaf blade and petiole (Figures 4A-D).

Because petiole hyponastic responses occur in the timeframe of hours and days, we next conducted long-term imaging (up to 8 hours) to capture long-term, sustained Ca^2+^ dynamics in leaves. We observed petiole-localized Ca^2+^ spiking in Col-0, many of which behaved as Ca^2+^ waves that propagated along a significant length of the petiole (Figure 4E; Video S5). The onset of Ca^2+^ waves preceded high humidity-induced hyponasty in the imaging chamber by 2.3 hours on average (Figure 4G). The *fer-4* mutant exhibited significantly fewer petiole-localized Ca^2+^ waves, but the spikes were longer in duration when they did occur (Figures 4F and 4H; Video S6; Figure S7). Altogether, these data suggest significant alterations in petiole-localized long-term Ca^2+^ signaling in the *fer-4* mutant that are associated with observed hyponastic and molecular phenotypes.

## DISCUSSION

In this study, we show that FERONIA is an important regulator of Arabidopsis high humidity responses. Whole-plant Ca²⁺ imaging showed that shifting plants from ambient to high humidity triggered cytosolic Ca²⁺ elevations, a response conserved across Arabidopsis, *Nicotiana benthamiana*, and *N. tabacum*, including petiole-localized Ca²⁺ waves preceding hyponasty. In the Arabidopsis *fer-4* mutant, these Ca²⁺ waves were altered, the early transcriptional signature was greatly diminished, including genes previously reported to contribute to humidity responses, and leaf hyponasty was impaired. These data are consistent with a model in which FERONIA is a critical receptor kinase that connects extracellular high humidity cues to petiole Ca²⁺ signaling, transcriptional reprogramming, and adaptive growth.

Our results position FERONIA transcriptionally upstream of a recently described high humidity signaling module. Hussain et al. 2024^<u>15</u>^ demonstrated that CNGC2/4-mediated Ca²⁺ influx activates CAMTA transcription factors, inducing ABA catabolism. However, in the *fer-4* mutant, *CNGC2* and *CYP707A3* were not induced despite normal *CAMTA* expression, suggesting that FERONIA regulates *CNGC2* and *CYP707A3* expression independently of and/or in parallel of the *CAMTA*-mediated mechanism. Jiang et al. 2024^<u>4</u>^ reported essential roles for ethylene in high humidity-mediated hyponasty. In this study, we found that petiole *ACS11* was rapidly induced in a FERONIA-dependent manner. Therefore, FERONIA may support local ethylene production during high humidity response. The reduced peak amplitude and altered spatial distribution of Ca²⁺ signals in the *fer-4* mutant is consistent with prior studies implicating FERONIA in Ca²⁺-dependent processes such as during pollen tube reception, root mechanosensing, and RALF peptide signaling^<u>38–40</u>^.

Our transcriptome analysis revealed rapid induction of genes involved in cell wall modification, including cellulose, hemicellulose, and especially pectin biosynthesis (*GAUTs*, *GALS*s, *GALT*s, *GAE1*) and modification (*OSU1*, *PMR6*, *PME*s, *PMEI*s, *PAE9*), pointing to cell wall dynamics as an important feature of the high humidity response. Pectin can absorb water, bind free Ca²⁺, and interact directly with FERONIA’s malectin-like ectodomain, with binding influenced by the degree of methylesterification^<u>22,28–30,41</u>^. Interestingly, our transcriptomic and biochemical data show that FERONIA is required for humidity-dependent changes in pectin methylesterification (Figures S1E and S4), suggesting a link FERONIA signaling to wall dynamics under high humidity.

FERONIA is a receptor for RALF peptides, yet exogenous RALF1 did not significantly affect humidity-dependent petiole elongation in our study, suggesting that FERONIA’s role in high humidity response is independent of RALF1. However, we cannot rule out the possibility that spraying RALF1 may not reach to the responsive petiole cell layers or reproduce the localized signaling gradients needed for asymmetric growth between the adaxial and abaxial sides. The Arabidopsis genome encodes over 30 distinct RALF peptides, many with specialized tissue-specific and stimulus-specific roles in FER signaling. It is formally possible that shoot humidity responses may operate through FERONIA sensing of a distinct RALF.

Whether FERONIA functions as a direct high humidity sensor itself, or serves primarily as a transducer of high humidity signaling, remains to be further delineated in future studies. Recent studies suggest that FERONIA-RALFs-PMEs module together acts as a cell wall integrity sensor^<u>41,42</u>^. It is therefore possible that high humidity alters cell wall structures, which are sensed directly by the FERONIA receptor. Alternatively, FERONIA may act downstream of indirect cues generated by high humidity transitions. Rapid increases in humidity are known to alter cellular turgor pressure^<u>43</u>^, which could be transduced via mechanosensitive signaling pathways^<u>44,45</u>^. However, we found that several mutants in putative mechanosensor genes responded normally to high humidity (Table S2), consistent with the reported response of the *msl10* mutant^<u>4</u>^. Additionally, high humidity potentially suppresses transpiration^<u>46</u>^, limiting gas diffusion and causing endogenous gases to accumulate^<u>4</u>^. Distinguishing these possibilities will require physical and biophysical measurements of plant cells under different humidity conditions. Nevertheless, our finding that FERONIA plays a critical role in multiple high humidity-induced responses, including genome-wide transcriptional changes, Ca²⁺ dynamics, petiole elongation, and hyponastic leaf movements, represents an important step in the understanding of how plants adapt to high humidity, one of the most important environmental conditions they encounter in nature.

### Limitations of the study

Our bulk RNA-seq time course cannot resolve transient or cell-type-specific changes or post-translational regulation. The lack of an effect of exogenous RALF1 does not rule out a role for RALFs in high humidity–induced hyponasty, as applied peptides may not reproduce the timing, dose, and local context of endogenous release. Whether high-humidity-induced changes in the turgor pressure or cell wall mechanics precedes FERONIA signaling remains to be investigated.

## RESOURCE AVAILABILITY

### Lead contact

Further information or requests for resources and reagents should be directed to and will be fulfilled by the lead contact, Sheng Yang He.

### Materials availability

All materials generated in this study are available upon request, subject to completion of MTA.

### Data and code availability

✰ All data reported in this paper will be shared by the lead contact upon request.
✰ RNA-Seq data have been deposited at GEO under accession GEO: GSE308941 and are publicly available as of the date of publication.
✰ No original code is reported in this paper.
✰ Additional information required to reanalyze the data reported in this paper is available from the lead contact upon request.

## ACKNOWLEDGEMENTS

This study was supported by funding from Chalermphrakiat Grant, Faculty of Medicine Siriraj Hospital, Mahidol University, Thailand, to C.O., United States Department of Agriculture Postdoctoral Fellowship 2020-10954 to R.H., Chinese Academy of Agricultural Sciences (Institute of Plant Protection) Postdoctoral Scholarship (2022M723452) to N.W., and Duke University to S.Y.H. S.Y.H. is an Investigator at Howard Hughes Medical Institute. We thank the excellent staff in the Duke University Phytotron and Greenhouse for plant growth support.

## AUTHOR CONTRIBUTIONS

S.Y.H., C.O., R.H., and K.N. conceptualized and designed this study; C.O., R.H., J.Z., S.P., N.W., C.K., and K.N. performed experiments and analyzed data; C.O., R. H., and S.Y.H. analyzed data; C.O., R.H., and S.Y.H. wrote the paper with all authors approved the final article.

## STAR METHODS

### KEY RESOURCES TABLE

| REAGENT or RESOURCE | SOURCE | IDENTIFIER |
| --- | --- | --- |
| **Antibodies** |  |  |
| Rat monoclonal JIM5 | University of Georgia, Athens, USA | Cat#JIM5 |
| Rat monoclonal JIM7 | University of Georgia, Athens, USA | Cat#JIM7 |
| Rat monoclonal LM19 | Kerafast | Cat#ELD001 |
| Rat monoclonal LM20 | Kerafast | Cat#ELD003 |
| Horseradish peroxidase (HRP)-conjugated Goat anti-rat IgG | Thermo Scientific | Cat#31470 |
| **Chemicals, peptides, and recombinant proteins** |  |  |
| Chemically synthesized RALF1 peptide<br>Sequence:ATTKYISYQSLKRNSVPCSRRGASYYN<br>CQNGAQANPYSRGCSKIARCRS | GenScript | N/A |
| **Critical commercial assays** |  |  |
| RNeasy Plant Mini Kit | QIAGEN | Cat#74904 |
| RNase-Free DNase set | QIAGEN | Cat#79254 |
| Plant RNA Pico Assay | Agilent | Cat#5067-1513 |
| SuperSignal™ West Dura Extended Duration Substrate | Thermo Scientific | Cat#34075 |
| **Deposited data** |  |  |
| Raw and analyzed data | This paper | GEO: GSE308941 |
| **Experimental models: Organisms/strains** |  |  |
| *Arabidopsis: fer-4* | Arabidopsis Biological Resource Center (ABRC) | CS69044 |
| *Arabidopsis: llg1-1* | ABRC | CS66105 |
| *Arabidopsis: camta2/3* | ABRC | CS71604 |
| *Arabidopsis: siz1* | ABRC | CS862769 |
| *Arabidopsis: snc1* | Li, et al., 2001<sup>47</sup> | N/A |
| *Arabidopsis: UBQ10::R-GECO1*, Columbia-0 (Col-0) | Keinath, et al., 2015<sup>27</sup> | CS72772 |
| *Arabidopsis: UBQ10::R-GECO1, fer-4* | This study | N/A |
| *Arabidopsis: 35S::GCaMP3*, Col-0 | Toyota, et al., 2018<sup>48</sup> | N/A |
| *N. benthamiana: 35S::GCaMP3* | DeFalco, et al., 2017<sup>49</sup> | N/A |
| *N. tabacum*: 35S::GCaMP3 | DeFalco, et al., 2017<sup>49</sup> | N/A |
| Software and algorithms |  |  |
| ImageJ | Schindelin et al., 2012<sup>50</sup> | https://imagej.nih.gov/ij/ |
| STAR aligner | Dobin et al., 2013<sup>51</sup> | https://github.com/alexdobin/STAR |
| Subread | Liao et al., 2013<sup>52</sup> | https://subread.sourceforge.net |
| DESeq2 | Love et al., 2014<sup>53</sup> | https://github.com/thelovelab/DESeq2 |
| dplyr | Wickham et al., 2014<sup>54</sup> | https://dplyr.tidyverse.org |
| ggplot2 | Wickham et al., 2016<sup>55</sup> | https://ggplot2.tidyverse.org |
| pheatmap | Kolde et al., 2010<sup>56</sup> | https://github.com/raivokolde/pheatmap |
| SciPy version 1.0 | Virtanen et al., 2020<sup>57</sup> | https://docs.scipy.org/doc/scipy/index.html |
| μManager version 2.0 software | Edelstein et al., 2014<sup>58</sup> | https://micro-manager.org/Version_2.0 |
| Matplotlib | Hunter et al., 2007<sup>59</sup> | https://matplotlib.org |
| GraphPad Prism version 8 | GraphPad Software, Boston, MA | https://www.graphpad.com/ |

### EXPERIMENTAL MODEL AND STUDY PARTICIPANT DETAILS

#### Plant Materials and Growth Conditio*ns*

*Arabidopsis thaliana*, *Nicotiana benthamiana,* and *Nicotiana tabacum* seeds were stratified at 4°C for 2-4 days. Stratified seeds were grown on Arabidopsis mix soil (1:1:1 SureMix:vermiculite:perilite) in air-circulating growth chambers at ∼60% relative humidity, a temperature range of 20°C to 22°C, and light intensity of 100 µmol m^-2^ s^-1^ with a 12-hour light, 12-hour dark photoperiod for 4 to 4.5 weeks. T-DNA insertion mutants, listed in Table S2, were obtained from the Arabidopsis Biological Resource Center (ABRC) at Ohio State University. The *UBQ10::R-GECO1* biosensor line was crossed with *fer-4* lines and F_3_ homozygous lines were used for Ca^2+^ imaging experiments.

### METHOD DETAILS

#### High Humidity Treatment

The same growth chamber was used for ambient and high humidity conditions. Four- to four-and-a-half-week-old plants were placed in a plastic flat with a clear dome and a humidity sensor (SensorPush, #HT.w) to monitor humidity fluctuations during the day and night times. Fine-mist water spray applied beforehand to create high humidity condition inside the flat (at least 95% RH). Plants under high humidity were subjected for imaging after the treatments. To measure the petiole length, three to four fully expanded leaves at similar ages or middle whorls of the rosette were detached with fine forceps near the center of rosettes and imaged. Leaf petiole lengths were measured using Fiji and % relative change in length was calculated as ((L_HH_ – L_Amb_) / L_Amb_) x 100) where L_HH_ and L_Amb_ represent the length measurements under high humidity (HH) and ambient (Amb) conditions, respectively. To assess hyponastic movement, time-lapse videos captured side-viewed plants exposed to >95% RH high humidity for 8-12 hours. Fully expanded leaves (2-3 leaves per plant) were scored every two hours, and the percentages of upward-moving leaves relative to the total leaf number were calculated.

#### RNA Extraction and Gene Expression Analyses

RNA was extracted from flash-frozen Arabidopsis leaf or petiole tissue using RNeasy Plant Mini Kit (QIAGEN, #74904) and DNA contaminants were removed using RNase-Free DNase set (QIAGEN, #79254) following the manufacturer’s protocols. A minimum of three biological replicates with RNA integrity scores at least 7 using the Agilent Bioanalyzer 2100 and Plant RNA Pico Assay (#5067-1513) were chosen for library preparation and sequenced using Illumina NovaSeq 6000 in a 150bp PE format at the Michigan State University Genomics Core Facility. Data was deposited in the Gene Expression Omnibus with the GEO accession # GSE308941. Reads were mapped and counted by STAR aligner and Subread. Differential expression analysis and data visualization were accomplished via R packages, including DESeq2, dplyr, ggplot2, and pheatmap.

#### Whole Plant Ca^2+^ Imaging and Analysis

Soil-grown *35S::GCaMP3* tobacco (4 to 5 weeks old), *35S::GCaMP3* Arabidopsis (3 weeks old), or *UBQ10::R-GECO1* Arabidopsis (4 weeks old) were imaged in a custom Alligator Luminescence System (Cairn). R-GECO1 lines were used for imaging 4-week-old plants. The chamber was equipped with an iXon 888 EMCCD camera (Oxford Instruments, Andor Technology), and a custom Optospin emission filter switching unit (Cairn). A pE-4000 (CoolLED) unit was connected to a fiberoptic cable with 4-splitter terminal mounted leads for excitation illumination of plant samples. The pE-4000 and Optospin units were equipped with 25mm diameter mounted optical filters (Chroma Technology) for imaging GCaMP3 (excitation = 470nm; excitation filter:ET470/40x; emission filter:ET535/30m)) and R-GECO1 (excitation = 550nm; excitation filter:550ET560/25x; emission filter:ET620nm/60m. For long-term imaging, the top surface of the chamber was fitted with white LEDs (∼100umol m-2 s-1), (9 minutes on, 1 minute off for fluorescence acquisition), to limit the impact of darkness on the plant physiology. Humidity was set to ∼60% RH for ambient imaging experiments and to >95% RH for high humidity imaging experiments. Image acquisition was controlled via ImageJ using µManager2.0 software.

Image analysis was performed using FIJI. Briefly, image stacks were auto-thresholded using the Default settings in the Thresholder tool to set background pixel values to NaN. The polygon tool was used to manually define region of interests (ROI)s. All data presented was normalized using the expression (F-F0)/F0, where F is the average intensity value for a ROI at a specific time point, and F0 is the initial average value for the same ROI. were defined as signal elevations exhibiting a signal-to-noise ratio of [(F-F0)/F0]>0.15. Petiole-localized Ca^2+^ spikes were identified using the find_peaks() function (12) from SciPy^<u>57</u>^, with thresholds for prominence ((F-F0)/F0 ≥ 0.15) and minimum peak distance (≥ 20 time points). Kymograms were created using FIJI’s kymogram tool by drawing an 11-pixel wide line from the base of the petiole to the leaf tip. The “fire” LUT was used for displaying Ca^2+^ levels in all GCaMP3 and R-GECO1 images. Plots and graphs for analysis were created using GraphPad Prism or the matplotlib library in Python.

#### Pectin Methylesterification Assay

Four- to five-week-old plants were subjected to either ambient (∼60%) or high humidity (RH >95%) for up to 6 days. Fully expanded leaves from the middle whorl were collected. Four mm leaf discs were taken from lamina tissue, avoiding major veins. Tissue fresh weight (FW) was recorded prior to cell wall extraction. For cell wall extraction, leaf discs (0.005–0.010 g FW) were homogenized with ceramic beads in extraction buffer (50 mM CDTA, 50 mM ammonium oxalate, 50 mM ammonium acetate, pH 5.5; 150 µL per 0.005 g FW) using a GenoGrinder (2 × 30 s, 1500 rpm). Extracts were mixed with 3 M sodium acetate and ethanol (5:1:5, v/v/v), precipitated at –80 °C for 1 h, and centrifuged at maximum speed for 20 min. Pellets were washed twice with 75% ethanol, air-dried, and resuspended in water (50 µL per 0.005 g FW). For immuno-dot blot assay, nitrocellulose membranes were pre-wetted with TBS-T (20 mM Tris, 150 mM NaCl, 0.1% Tween-20) and mounted on a vacuum blotting apparatus. Two microliters of resuspended cell wall extract were spotted and air-dried under vacuum. Membranes were blocked with 10% non-fat milk in TBS-T for 1 h at room temperature and incubated overnight at 4°C with the primary monoclonal antibodies JIM5 and LM19 (de-esterified homogalacturonan) or JIM7 and LM20 (methyl-esterified homogalacturonan) (all at 1:1000 dilution in 5% milk/TBS-T). After washing, membranes were probed with an HRP-conjugated anti-rat IgG secondary antibody (1:5000, Thermo) for 45–60 min at room temperature. Following additional washes, signals were detected using a chemiluminescent substrate (SuperSignal™ Dura, Thermo).

### QUANTIFICATION AND STATISTICAL ANALYSIS

Statistical tests, exact n values, what n represents, and definitions of center and dispersion are reported in the figure legends. Data are presented as mean ± SEM unless otherwise stated. RNA-seq boxplots show the median and interquartile range. Petiole elongation and leaf movement data (Figures 2 and S6) were analyzed by two-tailed Mann-Whitney U test. Ca²⁺ imaging parameters were analyzed via two way ANOVA followed by Tukey’s post hoc test (Figure 4B, D) or were analyzed by Student’s t-test (Figure 4 F-I). RALF1 treatment effects (Figure S2C) were analyzed by Kruskal-Wallis test, and pectin de-esterification ratios (Figure S1E) by two-way ANOVA. Non-parametric tests were used so that normally distributed data did not need to be assumed. Significance was defined as p < 0.05. For RNA-seq, differentially expressed genes were identified with DESeq2 using a Benjamini-Hochberg adjusted p value (padj) < 0.05, with no fold-change threshold applied. Petiole Ca²⁺ spikes were identified with the SciPy find_peaks() function using a prominence threshold of (F−F₀)/F₀ ≥ 0.15 and a minimum peak distance of 20 time points. Statistical analyses were performed in GraphPad Prism 8 and Python (SciPy).

## SUPPLEMENTAL INFORMATION

**Video S1 (separate file), related to Figure 2**. Time-lapse video of upward leaf movements in Col-0, *fer-4* and *llg1-1* plants under high humidity (>95% RH) conditions for 8 hours.

**Video S2 (separate file), related to Figure 5**. High humidity-dependent Ca^2+^ signaling in 3-week-old *A. thaliana* plant expressing GCaMP3. Plant was shifted from ∼60% RH to >95% RH at time = 0 minutes.

**Video S3 (separate file), related to Figure 5**. High humidity-dependent Ca^2+^ signaling in 5-week-old *N. benthamiana* plant expressing GCaMP3. Plant was shifted from ∼60% RH to >95% RH at time = 0 minutes.

**Video S4 (separate file), related to Figure 5**. High humidity-dependent Ca^2+^ signaling in 5-week-old *N. tabacum* plant expressing GCaMP3. Plant was shifted from ∼60% RH to >95% RH at time = 0 minutes.

**Video S5 (separate file), related to Figure 4**. High humidity-dependent Ca^2+^ signaling in 4-week-old *A. thaliana* Col-0 wildtype leaf and petiole expressing R-GECO1. Plants were shifted from ∼60% RH to >95% RH at time = 0 minutes.

**Video S6 (separate file), related to Figure 4**. High humidity-dependent Ca^2+^ signaling in 4-week-old *A. thaliana fer-4* leaf and petiole expressing R-GECO1. Plants were shifted from ∼60% RH to >95% RH at time = 0 minutes.

**Table S1 (separate file), related to Figure 1**. Lists of DEGs responsive to high humidity in Col-0 leaves at 15 min, 60 min and 24 hours after high humidity (>95% RH) treatment.

**Table S2 (separate file).** Lists of T-DNA mutant lines used in this study.

