## Supplemental Information for "The FERONIA receptor kinase is required for high humidity responses in Arabidopsis"

Supplemental figures 1-7, with titles and legends

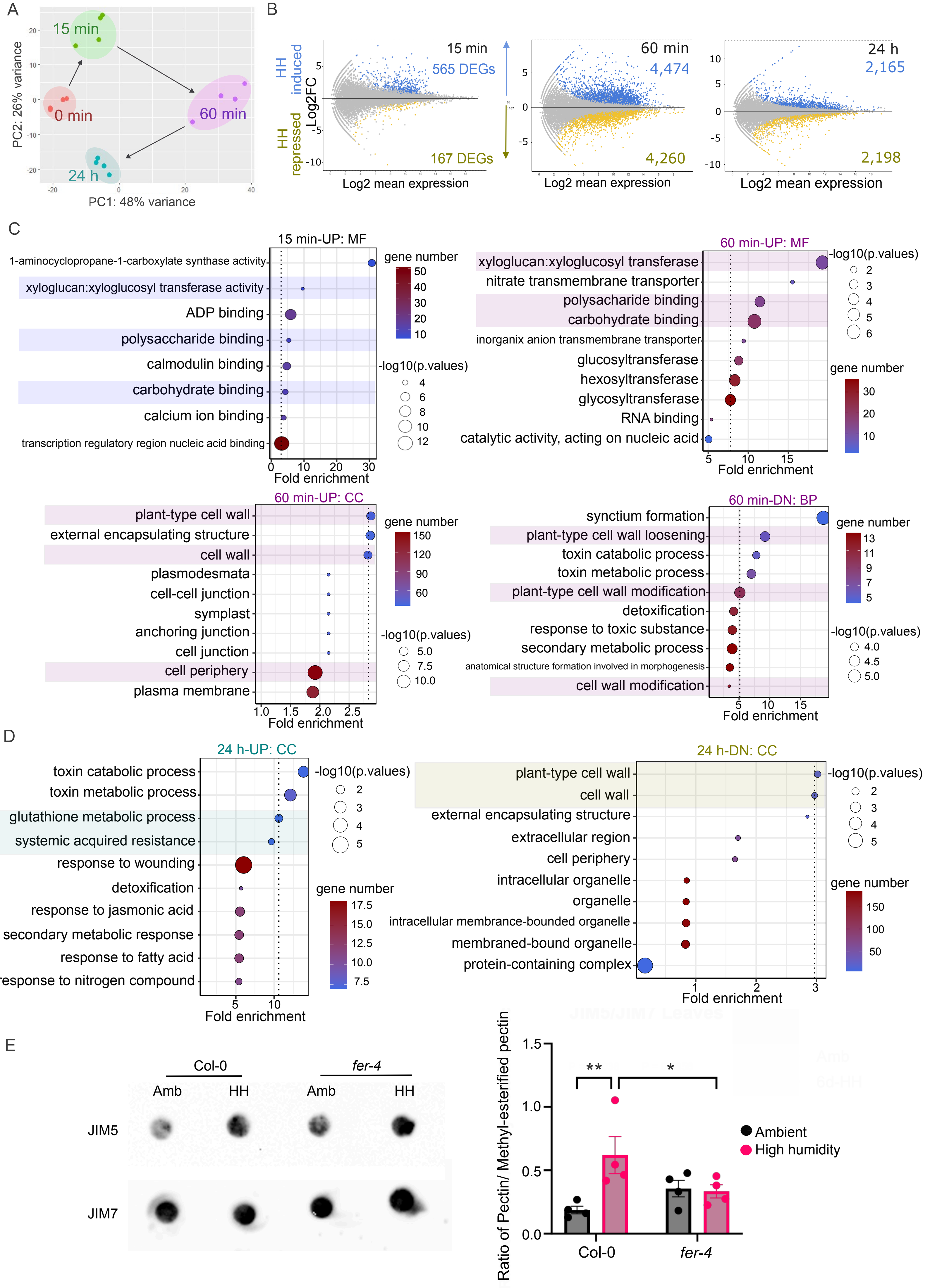
A
PC2: 26% variance
PC1: 48% variance
15 min
0 min
60 min
24 h
B
HH induced
HH repressed
Log2FC
15 min
565 DEGs
167 DEGs
Log2 mean expression
60 min
4,474
4,260
Log2 mean expression
24 h
2,165
2,198
Log2 mean expression
C
15 min-UP: MF
1-aminocyclopropane-1-carboxylate synthase activity
xyloglucan:xyloglucosyl transferase activity
ADP binding
polysaccharide binding
calmodulin binding
carbohydrate binding
calcium ion binding
transcription regulatory region nucleic acid binding
Fold enrichment
gene number
-log10(p.values)
60 min-UP: MF
xyloglucan:xyloglucosyl transferase
nitrate transmembrane transporter
polysacharide binding
carbohydrate binding
inorganix anion transmembrane transporter
glucosyltransferase
hexosyltransferase
glycosyltransferase
RNA binding
catalytic activity, acting on nucleic acid
Fold enrichment
60 min-UP: CC
plant-type cell wall
external encapsulating structure
cell wall
plasmodesmata
cell-cell junction
symplast
anchoring junction
cell junction
cell periphery
plasma membrane
Fold enrichment
60 min-DN: BP
synctium formation
plant-type cell wall loosening
toxin catabolic process
toxin metabolic process
plant-type cell wall modification
detoxification
response to toxic substance
secondary metabolic process
anatomical structure formation involved in morphogenesis
cell wall modification
Fold enrichment
D
24 h-UP: CC
toxin catabolic process
toxin metabolic process
glutathione metabolic process
systemic acquired resistance
response to wounding
detoxification
response to jasmonic acid
secondary metabolic response
response to fatty acid
response to nitrogen compound
Fold enrichment
24 h-DN: CC
plant-type cell wall
cell wall
external encapsulating structure
extracellular region
cell periphery
intracellular organelle
organelle
intracellular membrance-bounded organelle
membraned-bound organelle
protein-containing complex
Fold enrichment
E
Col-0
fer-4
Amb
HH
Amb
HH
JIM5
JIM7
Ratio of Pectin/ Methyl-esterified pectin
Ambient
High humidity
Col-0
fer-4
**
*

**Figure S1. Whole-leaf transcriptome analysis of Col-0 exposed to high humidity, related to Figure 1. (**A) Principal component analysis (PCA) plot displays the variation among four different timepoints (n=4). (B) MA plots show DEGs at three timepoints with the number of high humidity-induced and –repressed DEGs at each timepoints (Red dots indicate padj < 0.05). Numbers of up- and down-regulated DEGs at each timepoint are indicated in boxes (green and red, respectively; padj<0.05). (C and D) Enriched GO terms of high humidity-responsive DEGs at the 15-, 60-minute (C) and 24-hour (D) timepoints. Type of GO terms- biological process (BP), molecular function (MF) and cellular component (CC)- are indicated. The enrichment of high humidity-induced DEGs is indicated in blue and yellow represents high humidity-repressed DEGs. (E) Immunoblots of pectin methyl-esterification status in the leaves of Col-0 and the *fer-4* mutant with a quantification plot of de-esterified to methyl-esterified pectin ratios under ambient (black) and high humidity (HH; pink) conditions for 6 days. Statistical significance was assessed using Two-way ANOVA with Tukey's test. P values were indicated (*p < 0.05 and **p < 0.01). Bars represent mean ± S.E.M; n=4.

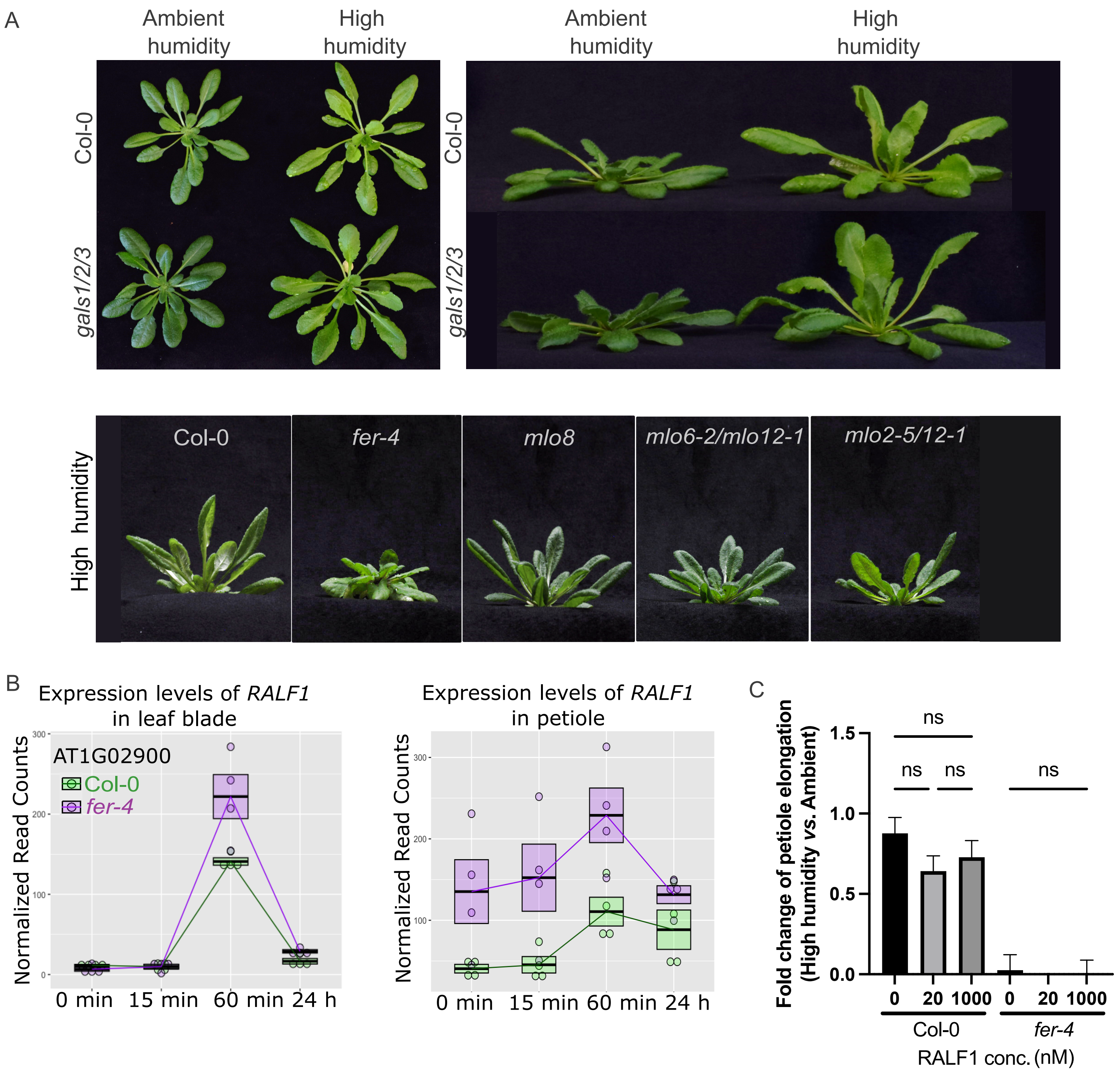

A
Ambient humidity
High humidity
Ambient humidity
High humidity
Col-0
gals1/2/3
Col-0
gals1/2/3
High humidity
Col-0
fer-4
mlo8
mlo6-2/mlo12-1
mlo2-5/12-1
B
Expression levels of RALF1 in leaf blade
AT1G02900
Col-0
fer-4
Normalized Read Counts
0 min 15 min 60 min 24 h
Expression levels of RALF1 in petiole
Normalized Read Counts
0 min 15 min 60 min 24 h
C
Fold change of petiole elongation (High humidity vs. Ambient)
ns
ns
ns
ns
1.5
1.0
0.5
0.0
0 20 1000 0 20 1000
Col-0
fer-4
RALF1 conc. (nM)

**Figure S2. Petiole elongation and upward leaf movement of Arabidopsis mutants, related to Figure 2.** (A) Pictures depicted petiole elongation and upward leaf movement under high humidity in Col-0 and the *gals1/2/3* triple mutant. Top panel: Top-viewed and side-viewed pictures. Bottom panel: Pictures depicted upward leaf positions under high humidity in Col-0, *fer-4, mildew resistance locus O (mlo) mlo-8, mlo6-2/mlo12-1,* and *mlo2-5/mlo12-1* mutants. (B) Effects of ectopic application of RALF1 peptide (1μM) on hyponasty in Col-0 plants under ambient (~60% RH) and high (>95% RH) humidity conditions. RNAseq plots of *RALF1* in leaf blades and petioles of Col-0 (green) and the *fer-4* mutants (purple) at different timepoints after exposure to high humidity. Boxes represent median ± quartiles 1 and 3; n=4. (C) Quantitation of petiole length elongation (Fold change) in Col-0 plants under 95% RH for 6 days compared to that in Col-0 plants under ~60% RH controls (Ambient RH). Col-0 and *fer-4* plants were pre-treated with water (control) or with 20 nM and 1 μM synthetic RALF1 peptide under ambient and high humidity conditions. Statistical significance was assessed using Kruskal-Wallis test and p value > 0.05 were indicated as non-significance (ns). Bars represent mean ± S.E.M; n>20.

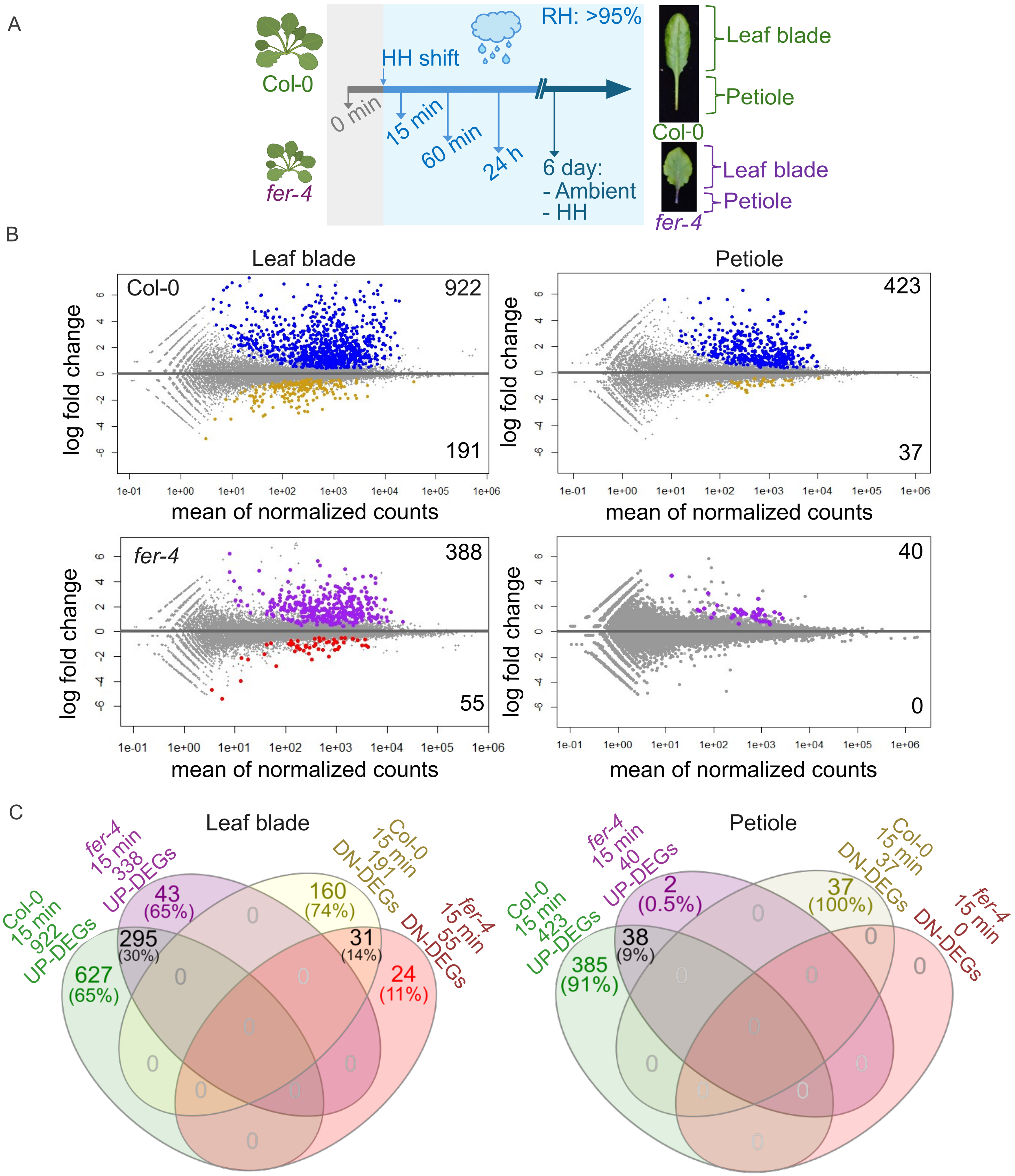

A
Col-0
fer-4
HH shift
RH: >95%
0 min
15 min
60 min
24 h
6 day:
- Ambient
- HH
Leaf blade
Petiole
Col-0
Leaf blade
Petiole
fer-4
B
Leaf blade
Petiole
Col-0
922
191
423
37
fer-4
388
55
40
0
log fold change
mean of normalized counts
1e-01 1e+00 1e+01 1e+02 1e+03 1e+04 1e+05 1e+06
C
Leaf blade
Col-0 15 min 922 UP-DEGs
fer-4 15 min 338 UP-DEGs
Col-0 15 min 191 DN-DEGs
fer-4 15 min 55 DN-DEGs
627 (65%)
43 (65%)
160 (74%)
24 (11%)
295 (30%)
31 (14%)
Petiole
Col-0 15 min 423 UP-DEGs
fer-4 15 min 40 UP-DEGs
Col-0 15 min 37 DN-DEGs
fer-4 15 min 0 DN-DEGs
385 (91%)
2 (0.5%)
37 (100%)
38 (9%)

**Figure S3. Leaf-blade and petiole transcriptome analysis of Col-0 and the *fer-4* mutant exposed to high humidity, related to Figure 3.** (A) A schematic diagram illustrates the leaf and petiole RNA-seq experimental designs for 4-week-old Col-0 and *fer-4* plants subjected to high humidity (HH; >95% RH) for 15 minutes, 60 minutes, 24 hours, and 6 days. Ambient (Amb; ~60% RH) and HH samples were collected at the 6-day timepoint for leaf blade and petiole tissues. (B) Minus-average (MA) plots show DEGs at the 15-minute timepoint in leaf blade and petiole tissues. The number of HH-induced and –repressed DEGs are indicated at the top- and bottom-right of each plot (Colors indicates padj < 0.05). Top-panels are Col-0, while bottom-panels are the *fer-4* mutant. (C) Venn diagrams of the up- and down-regulated DEGs between Col-0 and the *fer-4* mutant in leaf blades. Only upregulated DEGs are showed for petioles. Numbers of overlapped DEGs are indicated with percentages in parenthesis.

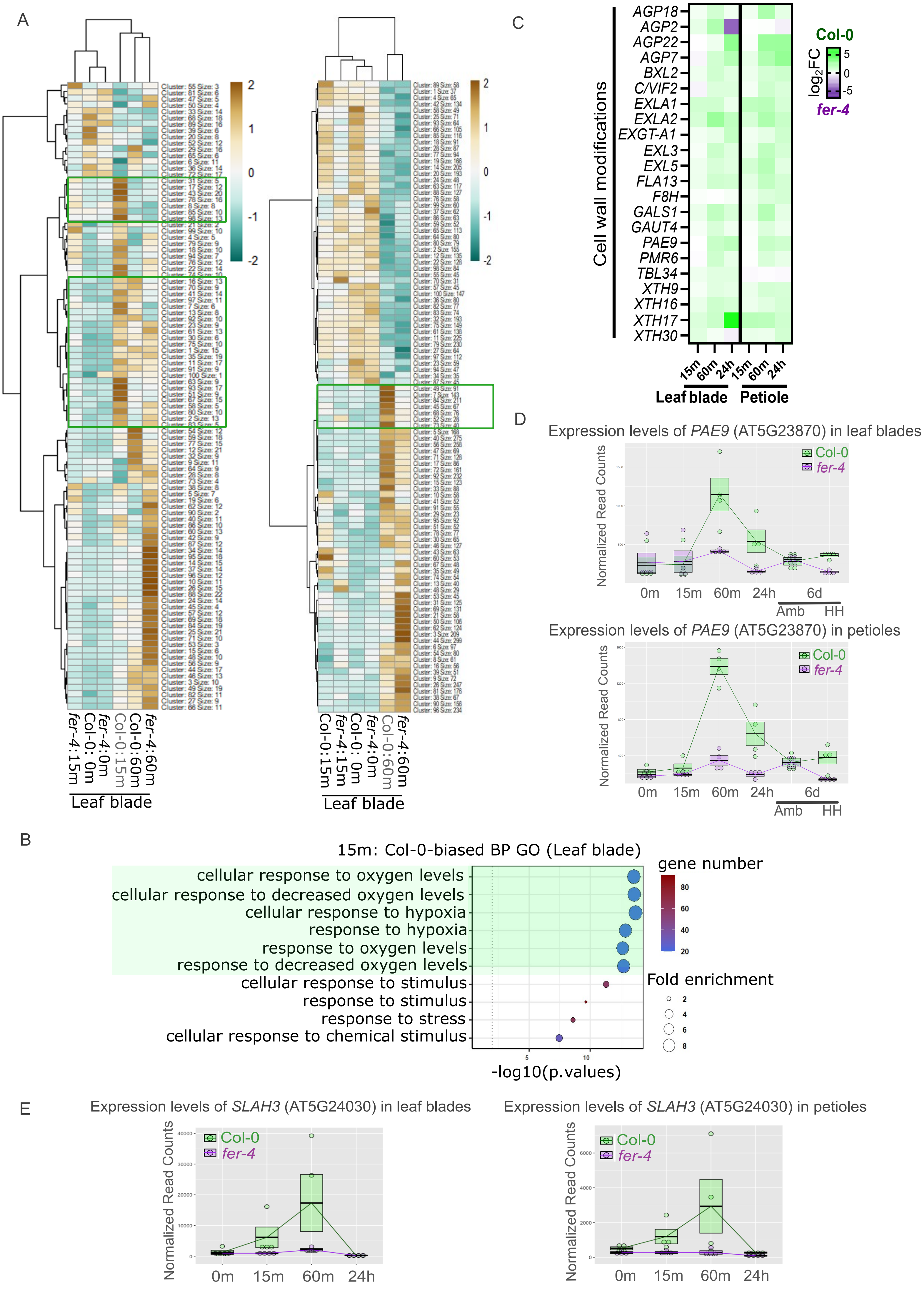
A
fer-4:15m
Col-0: 0m
fer-4:0m
Col-0:15m
Col-0:60m
fer-4:60m
Leaf blade
Col-0:15m
fer-4:15m
Col-0: 0m
fer-4:0m
Col-0:60m
fer-4:60m
Leaf blade
B
15m: Col-0-biased BP GO (Leaf blade)
cellular response to oxygen levels
cellular response to decreased oxygen levels
cellular response to hypoxia
response to hypoxia
response to oxygen levels
response to decreased oxygen levels
cellular response to stimulus
response to stimulus
response to stress
cellular response to chemical stimulus
-log10(p.values)
gene number
Fold enrichment
C
Cell wall modifications
AGP18
AGP2
AGP22
AGP7
BXL2
C/VIF2
EXLA1
EXLA2
EXGT-A1
EXL3
EXL5
FLA13
F8H
GALS1
GAUT4
PAE9
PMR6
TBL34
XTH9
XTH16
XTH17
XTH30
15m 60m 24h 15m 60m 24h
Leaf blade
Petiole
Col-0
log2FC
fer-4
D
Expression levels of PAE9 (AT5G23870) in leaf blades
Normalized Read Counts
Col-0
fer-4
0m 15m 60m 24h 6d
Amb HH
Expression levels of PAE9 (AT5G23870) in petioles
Normalized Read Counts
Col-0
fer-4
0m 15m 60m 24h 6d
Amb HH
E
Expression levels of SLAH3 (AT5G24030) in leaf blades
Normalized Read Counts
Col-0
fer-4
0m 15m 60m 24h
Expression levels of SLAH3 (AT5G24030) in petioles
Normalized Read Counts
Col-0
fer-4
0m 15m 60m 24h

**Figure S4. Transcriptome analysis of leaf blades of Col-0 and the *fer-4* mutant at the 15-min and 60-min timepoints after high humidity treatment, related to Figure 3.** (A) Clustering of high humidity-induced and –repressed DEGs identified in leaf blades of Col-0 and the *fer-4* mutant at the 15-minute and 60-minute timepoints. HH: High humidity; Amb: Ambient humidity. Groups of Col-0-biased DEGs at the 15-minute (leaf panel) and 60-minute (right panel) timepoints are labeled in a green box. Dark brown and dark green indicate Col-0-biased and *fer-4*-biased expression changes, respectively ($\log_2$ fold changes under high humidity *vs.* ambient humidity in Col-0 relative to *fer-4*). (B) Top enriched biological process gene ontology terms of Col-0-biased DEGs in leaf blade tissue ranked by adjusted p-values at the 15-minute timepoint. (C) A heatmap of selected cell wall-modifying DEGs in leaf blade and petiole tissues at three timepoints shows $\log_2$ fold changes (Col-0 *vs*. the *fer-4* mutant). Green indicates genes up-regulated in Col-0 relative to the *fer-4* mutant, while purple indicates genes with lower expression in Col-0 and higher expression in the *fer-4* mutant. (D and E) RNAseq plots of *PAE9* and *SLAH3* gene expression in leaf blade and petiole tissues of Col-0 (green) and the *fer*-4 mutant (purple) at different timepoints exposed to high humidity. Boxes represent median ± quartiles 1 and 3; n=4.

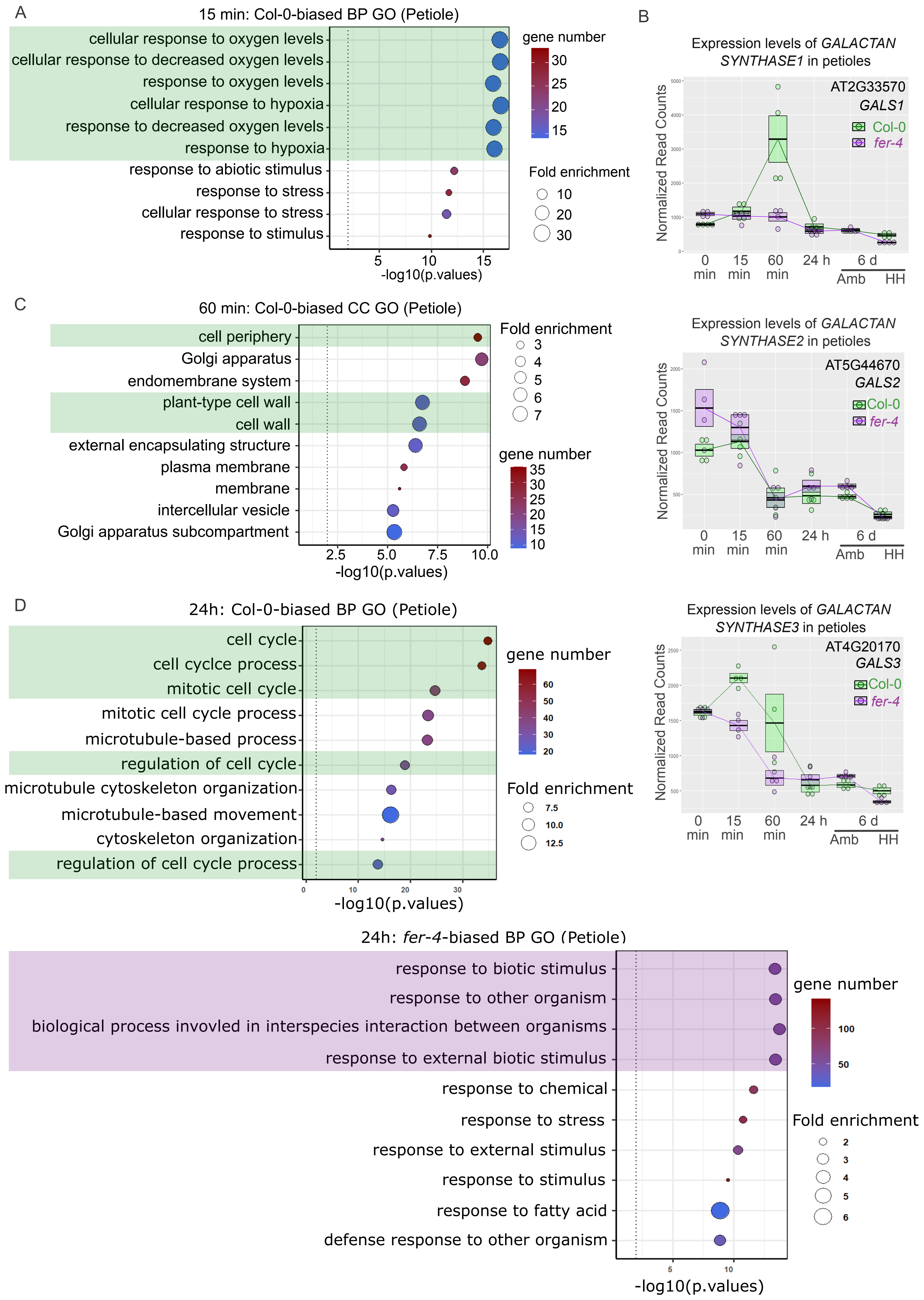

A
15 min: Col-0-biased BP GO (Petiole)
cellular response to oxygen levels
cellular response to decreased oxygen levels
response to oxygen levels
cellular response to hypoxia
response to decreased oxygen levels
response to hypoxia
response to abiotic stimulus
response to stress
cellular response to stress
response to stimulus
5
10
15
-log10(p.values)
gene number
30
25
20
15
Fold enrichment
10
20
30
B
Expression levels of GALACTAN SYNTHASE1 in petioles
Normalized Read Counts
AT2G33570
GALS1
Col-0
fer-4
0 min
15 min
60 min
24 h
6 d
Amb
HH
C
60 min: Col-0-biased CC GO (Petiole)
cell periphery
Golgi apparatus
endomembrane system
plant-type cell wall
cell wall
external encapsulating structure
plasma membrane
membrane
intercellular vesicle
Golgi apparatus subcompartment
2.5
5.0
7.5
10.0
-log10(p.values)
Fold enrichment
3
4
5
6
7
gene number
35
30
25
20
15
10
Expression levels of GALACTAN SYNTHASE2 in petioles
Normalized Read Counts
AT5G44670
GALS2
Col-0
fer-4
0 min
15 min
60 min
24 h
6 d
Amb
HH
D
24h: Col-0-biased BP GO (Petiole)
cell cycle
cell cyclce process
mitotic cell cycle
mitotic cell cycle process
microtubule-based process
regulation of cell cycle
microtubule cytoskeleton organization
microtubule-based movement
cytoskeleton organization
regulation of cell cycle process
0
10
20
30
-log10(p.values)
gene number
60
50
40
30
20
Fold enrichment
7.5
10.0
12.5
Expression levels of GALACTAN SYNTHASE3 in petioles
Normalized Read Counts
AT4G20170
GALS3
Col-0
fer-4
0 min
15 min
60 min
24 h
6 d
Amb
HH
24h: fer-4-biased BP GO (Petiole)
response to biotic stimulus
response to other organism
biological process invovled in interspecies interaction between organisms
response to external biotic stimulus
response to chemical
response to stress
response to external stimulus
response to stimulus
response to fatty acid
defense response to other organism
5
10
-log10(p.values)
gene number
100
50
Fold enrichment
2
3
4
5
6

**Figure S5. Transcriptome analysis of petioles of Col-0 and the *fer-4* mutant at the 15-minute, 60-minute, and 24-hour timepoints and immunoblot analysis of pectin modification status in response to high humidity, related to Figure 3.** (A) Top enriched biological process (BP) GO terms of Col-0-biased DEGs in petioles ranked by p-values at the 15-min timepoints. (B) RNAseq plots showing gene expression patterns of three *GALSs (GALS1,2* and *3)* in petioles of Col-0 and the *fer-4* mutant (green and purple, respectively) at 5 different timepoints. Box's represent median ± quartiles 1 and 3; n=4. (C) Top enriched cellular component (CC) GO terms of Col-0-biased DEGs in petioles ranked by p-values at the 60-minute timepoints. **(**D) Top enriched BP GO terms of Col-0-biased DEGs (top panel) and *fer-4*-biased DEGs (bottom panel) in petioles at the 24-hour timepoint, ranked by p-values.

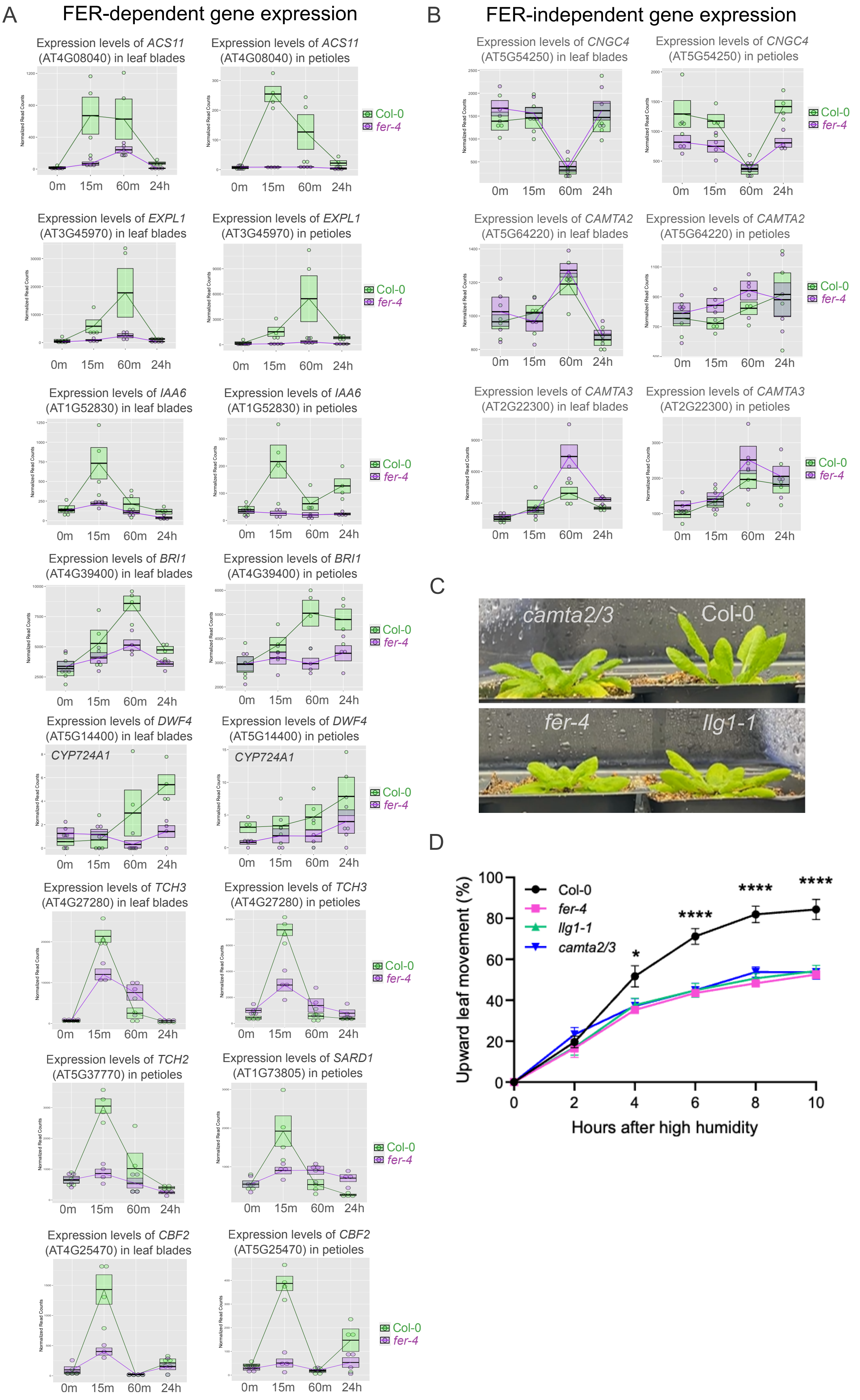
A
FER-dependent gene expression
Expression levels of ACS11 (AT4G08040) in leaf blades
Expression levels of ACS11 (AT4G08040) in petioles
Expression levels of EXPL1 (AT3G45970) in leaf blades
Expression levels of EXPL1 (AT3G45970) in petioles
Expression levels of IAA6 (AT1G52830) in leaf blades
Expression levels of IAA6 (AT1G52830) in petioles
Expression levels of BRI1 (AT4G39400) in leaf blades
Expression levels of BRI1 (AT4G39400) in petioles
Expression levels of DWF4 (AT5G14400) in leaf blades
Expression levels of DWF4 (AT5G14400) in petioles
CYP724A1
Expression levels of TCH3 (AT4G27280) in leaf blades
Expression levels of TCH3 (AT4G27280) in petioles
Expression levels of TCH2 (AT5G37770) in petioles
Expression levels of SARD1 (AT1G73805) in petioles
Expression levels of CBF2 (AT4G25470) in leaf blades
Expression levels of CBF2 (AT5G25470) in petioles
0m 15m 60m 24h
Col-0
fer-4
B
FER-independent gene expression
Expression levels of CNGC4 (AT5G54250) in leaf blades
Expression levels of CNGC4 (AT5G54250) in petioles
Expression levels of CAMTA2 (AT5G64220) in leaf blades
Expression levels of CAMTA2 (AT5G64220) in petioles
Expression levels of CAMTA3 (AT2G22300) in leaf blades
Expression levels of CAMTA3 (AT2G22300) in petioles
C
camta2/3
Col-0
fer-4
llg1-1
D
Upward leaf movement (%)
Hours after high humidity
Col-0
fer-4
llg1-1
camta2/3
*
****
****
****

**Figure S6. FERONIA-dependent and FERONIA-independent gene expression patterns in leaf blades and petioles of Col-0 and the *fer-4* mutant at the 15-minute, 60-minute, and 24-hour timepoints, related to Figures 2 and 3.** (A and B) RNAseq plots *1-AMINOCYCLOPROPANE-1-CARBOXYLATE SYNTHASE 11* (*ACS11)*, *EXPANSIN-LIKE FAMILY 1* (*EXPL1)*, *INDOLE-3-ACETIC ACID 6* (*IAA6)*, *BRASSIOSTEROID INSENSITIVE 1* (*BRI1)*, *DWARF4* (*DWF4)*, *CYCLIC NUCLEOTIDE GATED CHANNEL 4* (*CNGC4)*, *TOUCH-INDUCED* 2 (*TCH2*) and *TCH3, CALMODULIN-BINDING PROTEIN 60g (CBP60g), C-REPEAT BINDING FACTOR 2 (CBF2), SYSTEMIC ACQUIRED RESISTANCE DEFICIENT 1 (SARD1),* and *CALMODULIN-BINDING TRANSCRIPTION ACTIVATOR 2* (*CAMTA2)* and *CAMTA3* gene expression in two tissues of Col-0 and the *fer-4* mutant (green and purple, respectively) at 4 different timepoints. Boxes represent median ± quartiles 1 and 3; n=4. (C) Representative images of high humidity responses (hyponasty: petiole elongation and upward leaf movement) in Col-0, *fer-4*, *llg1-1* and *camta2/3* plants after 6 days of high humidity treatment. (D) Proportion of upward moving leaves under >95% RH high humidity condition during the first ten hours in Col-0 (black), the *fer-4* mutant (pink), the *llg1-1* mutant (green) and the *camta2/3* mutant (blue). Bars represent mean ± S.E.M; n=4. Mann-Whitney U test was used. P values were indicated (*$p < 0.05$ and ****$p < 0.0001$).

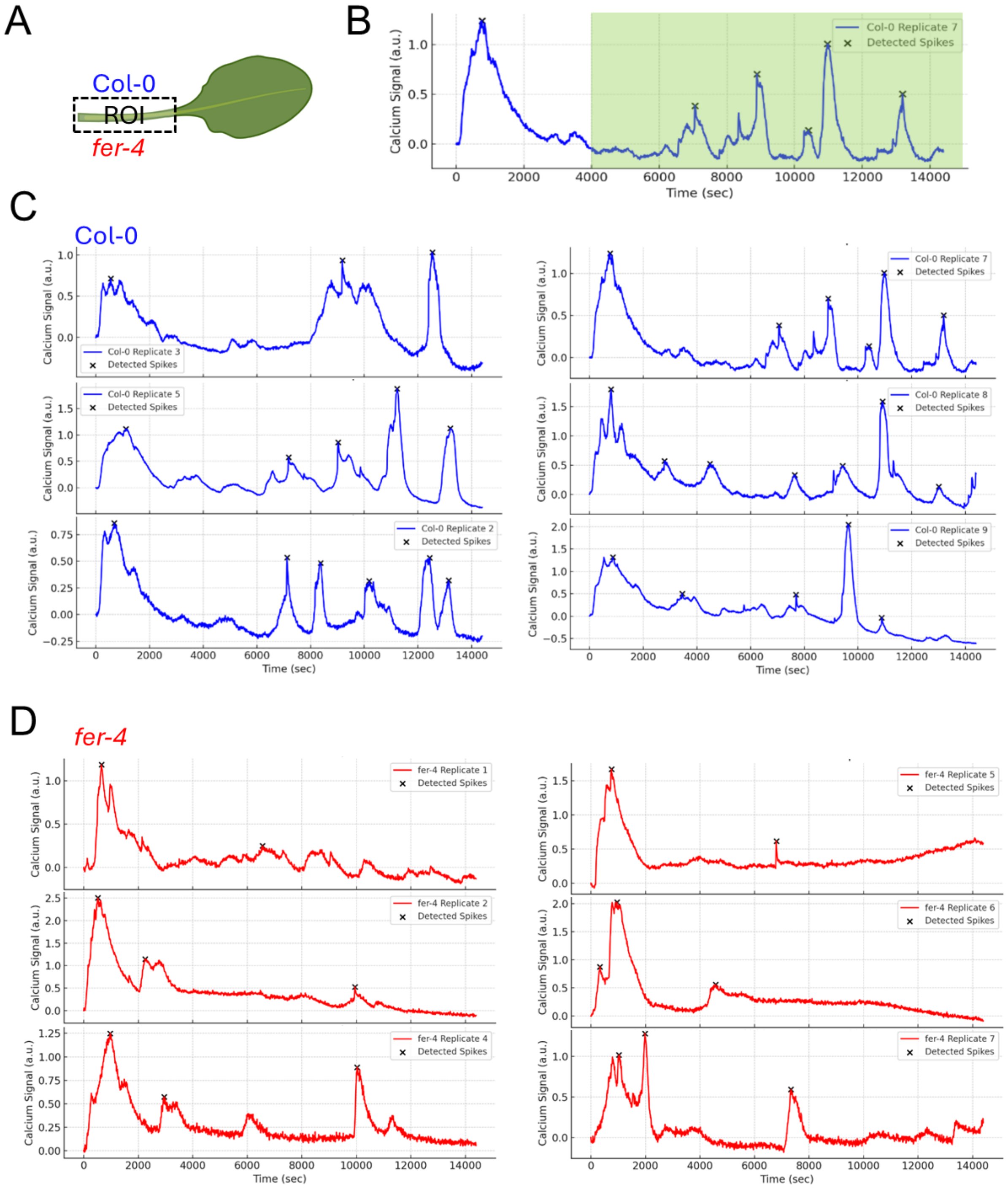

A
Col-0
ROI
fer-4
B
Col-0 Replicate 7
Detected Spikes
Calcium Signal (a.u.)
Time (sec)
C
Col-0
Col-0 Replicate 3
Col-0 Replicate 5
Col-0 Replicate 2
Col-0 Replicate 7
Col-0 Replicate 8
Col-0 Replicate 9
Detected Spikes
D
fer-4
fer-4 Replicate 1
fer-4 Replicate 2
fer-4 Replicate 4
fer-4 Replicate 5
fer-4 Replicate 6
fer-4 Replicate 7
Detected Spikes

**Figure S7. Petiole-localized $Ca^{2+}$ spiking in individual Col-0 and *fer-4* leaves in response to high humidity, related to Figure 4.** High humidity (>95% RH) elicits $Ca^{2+}$ spiking in Arabidopsis petioles. (A) A region of interest (ROI) in leaf petioles was selected and calcium spikes were assessed after 1.11 hours (4000 sec). (B) An example petiole $Ca^{2+}$ spiking trace indicating peak assessment time highlighted in green. $Ca^{2+}$ spike identification in (C) Col-0 and (D) *fer-4* petioles using find_peaks() function of SciPy. Peaks with a threshold signal >0.15 are marked with an “x.”
